# Engineering Microglial Cells to Promote Spinal Cord Injury Recovery

**DOI:** 10.1101/2024.07.09.602797

**Authors:** Qingsheng Zhou, Jianchao Liu, Qiongxuan Fang, Chunming Zhang, Wei Liu, Yifeng Sun

**Affiliations:** Spinal Surgery Department, YanTaiShan Hospital, Yantai, Shandong Province 264000, China; Department of Catheter room, Tianjin Chest Hospital, Tianjin, 300222, P. R. China.; MOE Key Laboratory of Cell Proliferation and Differentiation, School of Life Sciences, Peking University, Beijing 100871, China; Department of Orthopedic Surgery, The First Affiliated Hospital of Shandong First Medical University &Shandong Provincial Qianfoshan Hospital, Shandong Key Laboratory of Rheumatic Disease and Translational Medicine, Jinan, Shandong 250014, PR China; Orthopaedic and Sports Medicine Center, Beijing Tsinghua Changgung Hospital, Tsinghua University, China; Institute of Orthopedic and Sports Medicine of Tsinghua Medicine,Tsinghua University, China

**Keywords:** Microglia, Neuroinflammatory response, Microglial migration, TGFβ signal, Trem2, iPS-derived microglia

## Abstract

Spinal cord injury (SCI) can result in irreversible damage, leading to lifelong paralysis for affected individuals. Microglia’s dual impact on neuronal regeneration after SCI, driven by their distinct roles at different stages, merits further study. We conducted a bioinformatic analysis of single-cell transcriptomes (scRNA), spatial transcriptomic (ST) data, and bulk RNA-seq data from Gene Expression Omnibus (GEO) datasets. The data were processed using R packages such as “Seurat”, “DESeq2”,“limma” and “GSVA.” Additionally, we utilized Gene Set Enrichment Analysis (GSEA) and the Enrichr web servers. Analysis of single-cell data and spatial transcriptomics has revealed notable changes in the microglial cell landscape in SCI. These changes encompass the inhibition of innate microglial cells, while reactive microglial cells exhibit pronounced reactive hyperplasia. Moreover, the TGFβ signaling pathway plays a crucial role in regulating the migration of innate microglial cells to enhance SCI recovery. However, reactive microglial cells exhibiting high Trem2 expression contribute to the neuroinflammatory response and can effectively modulate neural cell death in SCI. In particular, inhibiting Trem2 in reactive microglial cells not only reduces inflammation but also mitigates spinal cord injury, and enhancing the TGFβ signaling pathway. What’s more, the use of iPSC-derived microglial cells, which have demonstrated their capacity to augment the potential for replacing the functions of naive microglial cells, iPSC-derived microglia have the potential to replace the functions of naive microglial cells, holds significant promise in addressing SCI. Therefore, we posit that the engineering of microglial cells to promote the SCI recovery. The approach of **i**nhibiting Trem2-mediated neuroinflammatory responses and transplanting iPSC-derived microglia with long-term TGFβ stimulation may offer potential improvements in SCI recovery.

## Introduction

Spinal cord injury (SCI) causes irreversible damage, leading to lifelong paralysis in patients (Simpson et al., 2012). In the spinal cord, neuron and glial cells interact actively, playing a vital role in neurofunction. Glial cells, comprising microglia, astrocytes, oligodendrocytes, and their progenitors, are the most abundant non-neuronal cell types (Gaudet & Fonken, 2018). When a spinal cord injury occurs, many glial cells at the lesion site become suppressed or activated, which further affects the inflammatory response and neuron regeneration after the injury (Orr & Gensel, 2018). Microglia is now recognized as an active participant in various pathological conditions in the central nervous system, such as trauma, stroke, and neurodegenerative disorders (Liu et al., 2011). Recent studies have revealed that microglia, much like a double-edged sword, can have both advantageous and disadvantageous effects on neuronal regeneration after SCI due to their distinct phenotypes and functions during different stages (Perry & O’Connor, 2010; Squarzoni et al., 2015; Trang et al., 2011; Ueno et al., 2013).

In the resting state, microglia play a crucial role in the development of the central nervous system (CNS) by contributing to the establishment of complex neuronal networks, controlling synaptic density, connectivity, and plasticity, and regulating the surrounding cellular milieu by secreting trophic factors, such as TGFβ and FGF, which promote the survival and regeneration of neurons (Utz et al., 2020). Furthermore, microglia exhibit a substantial presence of filamentous actin and display remarkable dynamism and motility in their resting state in vivo (Nimmerjahn et al., 2005). Upon injury, microglia migrate to the site of the lesion, where they not only begin to accumulate but also undergo morphological changes, potentially adopting pro-inflammatory characteristics as they become customized to the local environment (Kurpius et al., 2007). While activated microglia exacerbate neuronal injury through the production of pro-inflammatory cytokines, including Il-1α, TNF, and C1q, which in turn leads to secondary neuronal damage (Liddelow et al., 2017). Additionally, compelling evidence indicates that microglia actively engage in trauma-related injuries and neurodegenerative processes through a mechanism reliant on TREM2, thereby establishing a protective barrier that mitigates toxicity directed towards neighboring neurons (Yeh et al., 2017). However, it has been challenging to assign clear-cut roles to either cell subtype in vivo and to understand how microglia regulate a wide range of cellular and molecular responses following SCI (David & Kroner, 2011). Therefore, gaining a better understanding of the roles played by microglia in SCI is crucial for the development of effective therapies.

In this study, we conducted a integrated bioinformatic analysis that encompassed single-cell transcriptomes (scRNA), spatial transcriptomic data, and bulk RNA-seq data. Our investigation revealed that innate microglial cells exhibit heightened migratory activity, a phenomenon regulated by the TGFβ signaling pathway. However, during SCI, these cells undergo replacement by reactive microglia, which contribute to neuroinflammatory responses and exhibit up-regulated TREM2 expression. Inhibiting TREM2 has shown promise in reducing neuroinflammatory responses while enhancing the TGFβ signaling pathway. Additionally, the development of microglia derived from induced pluripotent stem cells (iPSC) represents an innovative approach, contributing to significant advancements in the field of SCI regeneration.

## Materials and method

### Single cell RNA sequencing (scRNA-seq) data processing

We utilized R statistical software (v. 4.1.2), the Seurat v4.0 toolkit, and MPLAB Harmony v1.0 to analyze scRNA-seq data. Public scRNA-seq data were obtained from the GSE189070, GSE150871, GSE198852 and GSE155499 datasets in the Gene Expression Omnibus (GEO). Low-quality cells (<3; features, <200; >10% mitochondrial genes) were filtered out, followed by initial normalization and removal of batch effects using LogNormalize (features = 3,000) and the Harmony function (max.iter.harmony = 20). Dimensionality reduction was performed using Uniform Manifold Approximation and Projection (UMAP).(Dai et al., 2022) To define the cell type with the marker genes identified by the Seurat package, we used PanglaoDB Augmented as described in our previous study (Sun et al., 2023).

### Spatial transcriptomics (ST) data analysis

We used Seurat to analyze the ST data, applying the same quality control and low-quality barcode filters used in scRNA-seq data analysis. In order to ensure that all sections were aligned to a common coordinate space, we devised a custom image analysis pipeline that encompasses pre-processing, registration, and the merging of histological images sourced from multiple sections. Cell types were identified in each subpopulation based on their known signatures and lineage markers, with innate- and adaptive-microglial cells identified by GSVA analysis, neurons by microtubule-associated protein 2 (Map2) and synaptosome associated protein 25 (Snap25).

### RNA Sequencing and Microarray data processing

To identify significantly differentially expressed genes (DEGs), we utilized a gene set comprising the expression profiles GSE218513, GSE196928, GSE89189 and GSE139192. A data matrix was obtained and used to annotate the probes into gene symbol sets. The Limma or Deseq2 package was employed for significance analysis, and the most significant changes in gene expression were determined based on false discovery rates and absolute fold change.

### Gene set enrichment analysis (GSEA)

GSEA is a computational method that determines whether an a priori defined set of genes shows statistically Significant .We assessed gene set enrichment using GSEA. BDNF regulation of GABA neurotransmission and Microglia Pathogen Phagocytosis Signaling pathway gene sets were curated from the Molecular Signatures Database (WikiPathway 2021 Human). Normalized enrichment scores (NES) >1 and p <0.05 reliably filtered significant pathways.

### Gene set variation analysis (GSVA)

Gene Set Variation Analysis (GSVA) is a nonparametric, unsupervised analysis method primarily employed for assessing the enrichment results of sequencing data with respect to gene sets. We established the molecular signatures of microglia subtypes depending on the significant marker genes, which were acquired by single cell analysis. We also calculated final scores for further analyses using GSVA (v.1.47.0), such as “Neuroinflammatory Response” and “Positive Regulation Of Neuron Death” biological process.

### Datasets and softwares

Enrichr (https://maayanlab.cloud/Enrichr/) is a powerful web server that hosts a diverse collection of datasets. For our analysis, we utilized several datasets, including WikiPathway signal pathway, GO Biological Process, and PanglaoDB Augmented. We considered statistical significance at a threshold of <0.05. Bioinformatic and statistical analyses were conducted using the R software (version 4.1.2) and GraphPad Prism 7.0 statistical software, respectively. A P-value of less than 0.05 was deemed to be statistically significant.

## Result

### Innate microglial cells are replaced by reactive microglia in SCI

To investigate the cellular landscape of SCI, we conducted a analysis of single-cell RNA sequencing data from uninjured and injured spinal cords at 7 and 14 days (GSE189070) (Hou et al., 2022). After performing quality control and batch correction, we obtained high-quality transcriptional data for 36834 cells, which were visualized using Uniform Manifold Approximation and Projection (UMAP). Using the generated markers (adj.P_Value<0.05; logFC>1), we identified a total of 15 clusters (Fig. 1A; Supplementary Table S1). Among the microglial cells analyzed, the pie plot and split UMAP cluster revealed that 97.5% of them exhibited innate microglia characteristics in the uninjured spinal cord. However, these numbers decreased to 3.1% at 7 days and 2.0% at 14 days post-injury, indicating a depletion in innate microglia at the lesion site following injury. Conversely, the proliferation of reactive microglial cells steadily increased at the lesion site after injury, rising from 2.5% in the uninjured state to 96.9% at 7 days and 98% at 14 days post-injury. (Fig. 1B, 1C). And the innate and reactive microglia displayed distinct gene signatures, potentially contributing to different biological processes. (Fig. 1D).

**Fig. 1.**
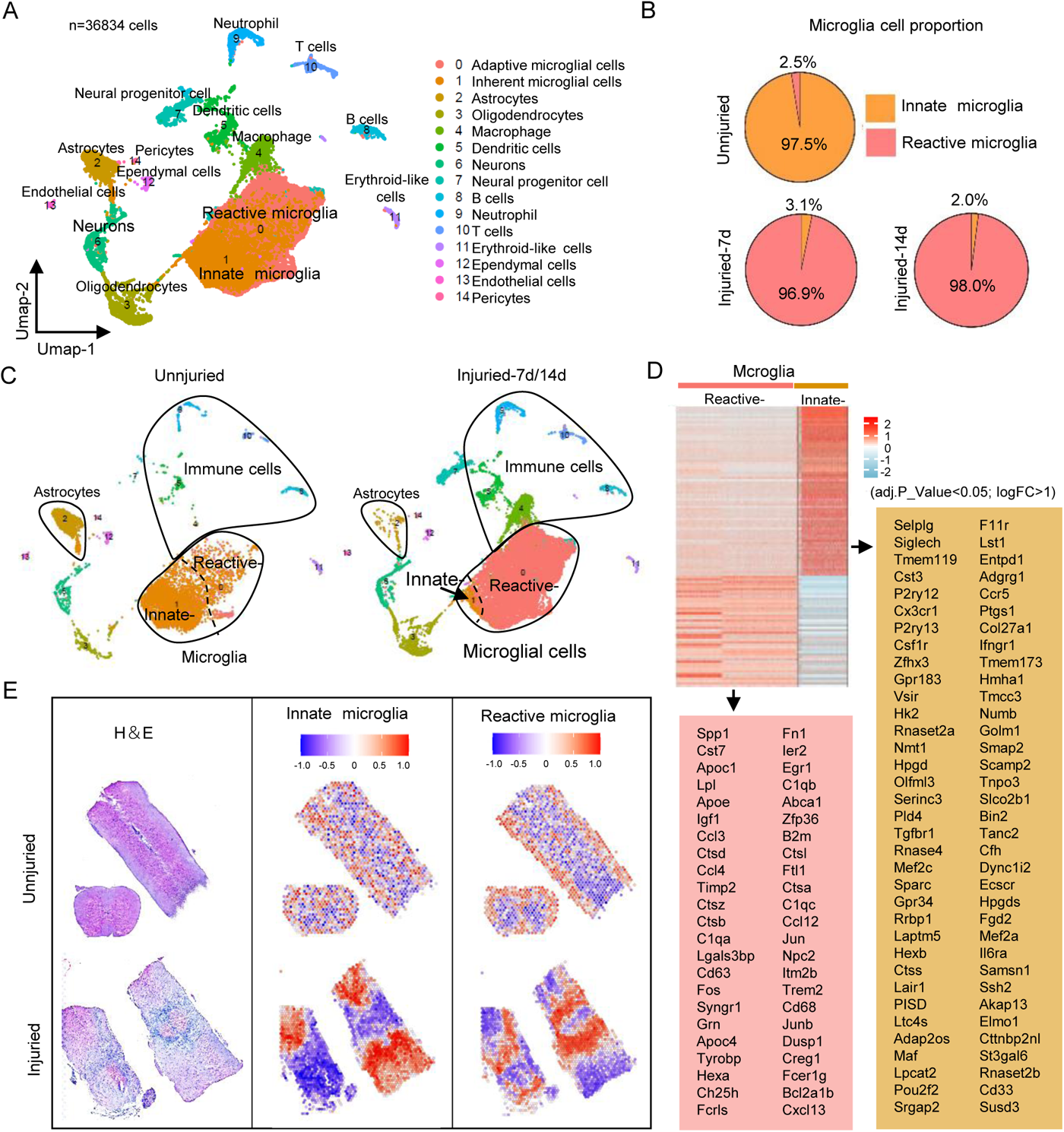
Innate microglial cells are replaced by reactive microglia in SCI. **(A)** Single-cell RNA sequencing (GSE189070) analysis was performed on both uninjured and injured spinal cords at 7 and 14 days post-injury. The resulting clusters were analyzed using the Uniform Manifold Approximation and Projection (UMAP) method, and different cell populations were identified and distinguished by color. **(B)** Pie plots was generated to visualize the changes in cell proportions for microglial cells. **(C)** Separated UMAP depending on uninjured and injured spinal cords. **(D)** Innate and reactive microglia gene signatures according to the differential gene expression analysis (DEGs) (adj.P_Value<0.05; logFC>1)). **(E)** The histological annotation (left), gene signature expression levels of innate (middle) and reactive (right) microglial cells in the spatial transcriptomics (ST) spots.

To confirm the aforementioned findings, we analyzed the published spatial transcriptomics (ST) dataset of contusion SCI (10 mice) or sham operation (2 mice) (GSE190910). We employed GSVA analysis to compute the scores for innate-and reactive-microglia using the previously described genes signatures, which were then mapped onto the histological images. Our analysis revealed a significant reduction in innate microglia and an augmented presence of reactive microglial cells (Fig. 1E). Hence, we conclude that the cascade of injury events in SCI significantly influences microglia, leading to the replacement of innate microglial cells by reactive microglia.

### TGFβ signaling pathway regulate innate microglial cell migration to improve SCI recovery

To investigate innate function of microglia, we utilized the innate microglial cell gene signatures identified from the scRNA data to conduct Gene Ontology Biological Process (BP) enrichment analysis. Our analysis revealed that, apart from the well-established ‘Negative regulation of lymphocyte activation,’ ‘Regulation of microglial cell migration’ was also significantly enriched, but it was reduced due to the decreased population of innate microglial cells during SCI (Fig. 2A). Within the realm of biological processes, 17 genes (Tgfbr1, Cx3cr1, Adgrg1, Akap13, F11r, Mef2c, P2ry12, Ccr5, Ctss, Hpgd, Hpgds, Laptm5, Lst1, Ptgs1, Samsn1, Tmem119, Zfhx3) actively participate in the top 10 processes. Notably, Tgfbr1 stands out as the most prevalent gene among this group of 17 (Fig. 2B). Meanwhile, in the STRING database, these 17 genes exhibit overlapping network modules, contributing to the formation of a protein-protein interaction (PPI) network. (Fig. 2C).

**Fig. 2.**
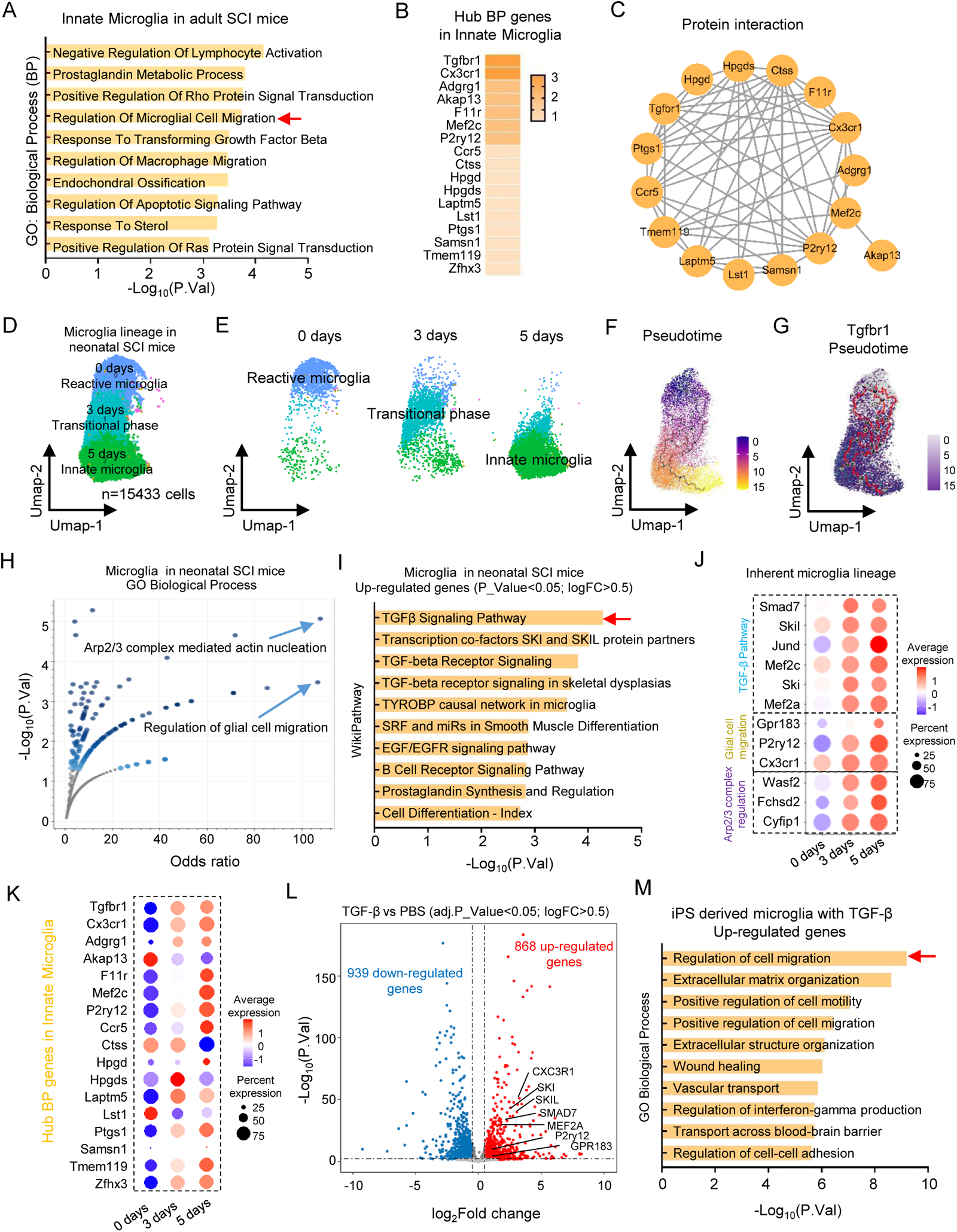
TGFβ signaling pathway regulate innate microglial cell migration to improve SCI recovery. (**A**) Enrichment analysis of Gene Ontology Biological Process (BP) using signatures from reactive microglial cells in single-cell data. (**B**) Hub genes within the top ten Biological Processes (BP) of reactive microglia. The intensity of color represents the frequency of observation. (**C**) Seventeen genes were utilized to create Protein-Protein Interaction (PPI) networks obtained from the STRING web interface and visualized using Cytoscape software. **(D)** Single cell analysis of Microglia lineage in neonatal SCI mice over the time course. The clusters were UMAP-analyzed, revealing distinct cell populations distinguished by color. **(E)** UMAP separation based on the time course, categorized by innate and reactive gene signatures. **(F)** Single-cell trajectory analysis conducted with Monocle3, cells color-coded according to their tissue origins and pseudotime. **(G)** UMAP visualization of cell dataset branch subsets, with cells color-coded according to Tgfbr1 expression. (H) Enriched BP in microglial cells from neonatal SCI mice, highlighting upregulated genes, visualized using Appyter (P<0.05; logFC>0.5). The volcano plot demonstrates the significance of each gene set from the selected library versus its odds ratio. (I) Pathway enrichment in microglial cells from neonatal SCI mice, showcasing upregulated genes, visualized using WikiPathway datasets. (J) Dot plot illustrating the expression of hub genes involved in the TGF-β signaling pathway, glial cell migration, and Arp2/3 complex-mediated actin nucleation within the innate microglia lineage. (K) Expression levels of 17 hub genes in innate microglia from neonatal SCI mice. (L) Volcano plot displaying significantly different genes (| log2FC | > 0.5; P < 0.05) between TGF-β and PBS-treated iPSC-derived microglia cells. (M) Enrichment analysis of Biological Processes using upregulated genes between TGF-β and PBS-treated iPSC-derived microglia cells.

Yi Li et al. have demonstrated that neonatal mice with spinal cord crush injuries can heal without scarring, allowing for the growth of long projecting axons through the lesion (Yi Li et al., 2020). In this study, we conducted an analysis of single-cell data obtained from microglial cells of neonatal mice under three different conditions (0, 3, and 5 days) within three distinct clusters (GSE150871) (Fig. 2D). In accordance with the expression changes in the gene signatures depicted in Supplementary Fig. S1, three distinct clusters were identified: Cluster 0 represented the immediate SCI with reactive microglia, Cluster 1 denoted the transitional phase at 3 days post-SCI, and Cluster 2 corresponded to 5 days post-SCI, where the microglia had already reverted to the innate state (Fig. 2E). We then conducted trajectory analysis to gain a deeper understanding of the transition of microglial cells, aligning perfectly with the timeline (Fig. 2F). These trajectory analyses provided enhanced clarity regarding gene dynamics along a specific pathway. Remarkably, we observed a significant increase in Tgfbr1 expression over the course of the timeline in neonatal SCI mice (Fig. 2G). The biological process “Glial cell migration”, and “Arp2/3 complex mediated actin nucleation”, as well as the TGF β signaling pathway were elevated in this neonatal SCI (Fig. 3H, 3I). Furthermore, the hub genes of TGFβ signaling pathway (Smad7, Skil, Jund, Mef2c, Ski, Mef2a), Arp2/3 complex mediated actin nucleation (Wasf2, Fchsd2, Cyfip1), and gial cell migration (Gpr183, P2ry12, Cx3cr1) were gradually increased with the development of neonatal SCI mice (Fig. 3J). Meanwhile, the 17 hub genes described above in innate microglial cell were also gradually increase with the timeline development (Fig. 3K). We then analyzed a comparative gene expression profiling analysis of RNA-seq data for iPSC derived microglia cells before and after TGFβ treatment, and identified 1807 DEGs (|log2FC| > 0.5; adj. P_val < 0.05), of which 868 were elevated in TGFβ treated iPSC-microglia (Fig. 3L) (GSE218513). Interestingly, the enrichment analysis revealed that TGFβ treatment significantly elevated microglial cell migration (Fig. 3M).

**Fig. 3.**
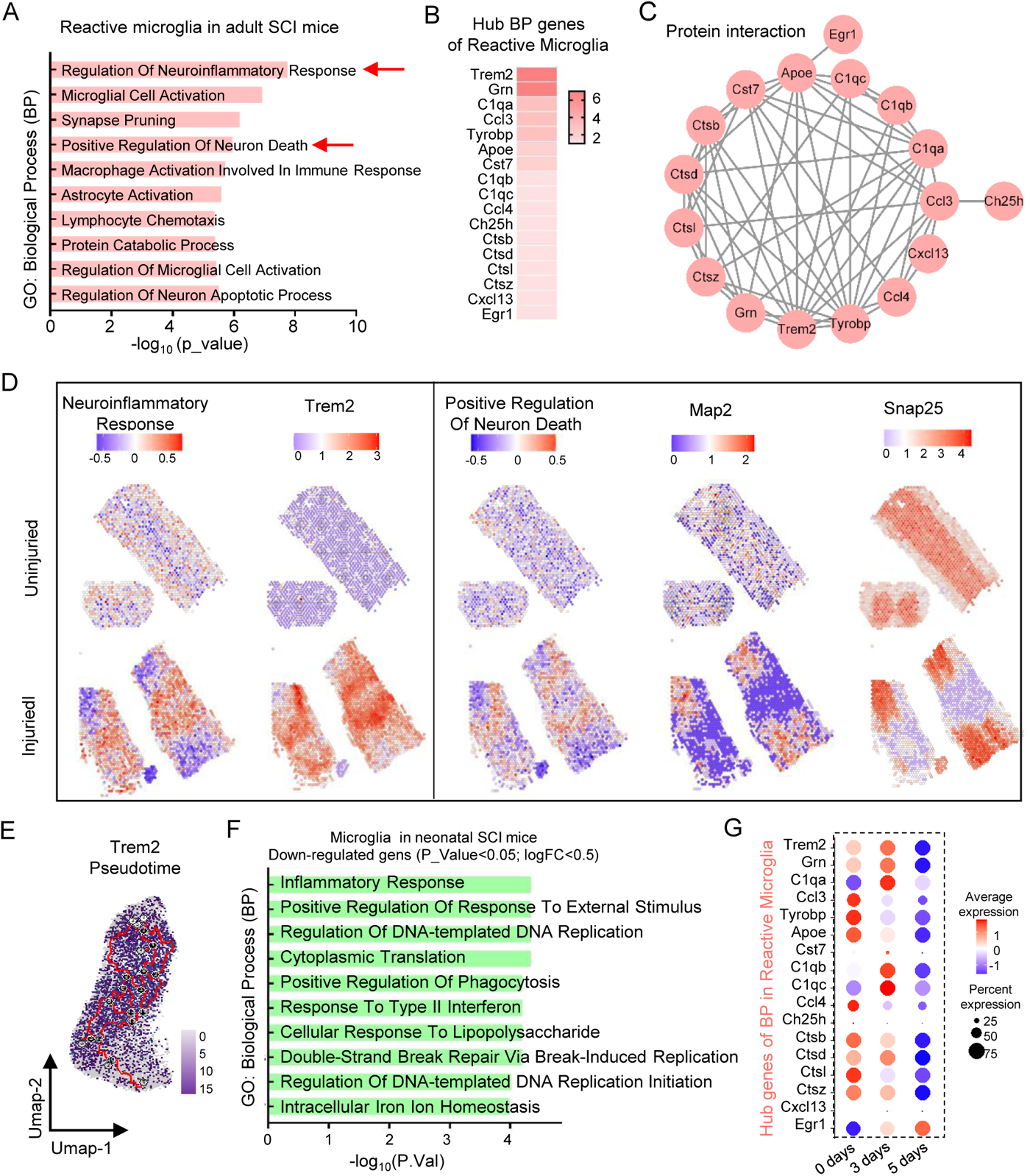
Reactive microglia contribute to the neuroinflammatory response and positive regulate the neural cell death in SCI. **(A)** Enrichment analysis of Gene ontology Biological Process (BP) using reactive microglial cell signatures in single cell data. **(B)** Hub genes of BP within reactive microglia’s top ten processes. The darker the color, the more frequently it is observed. **(C)** Seventeen genes were used to construct Protein-Protein Interaction (PPI) networks retrieved from the STRING web interface and visualized using Cytoscape software. **(D)** The gene signature scores for ‘Neuroinflammatory Response’ and ‘Positive Regulation Of Neuron Death,’ along with the standard Trem2 and neuron markers (Map2 and Snap25), were assessed in the spatial transcriptomics (ST) spots. **(E)** UMAP of cell data set branch subsets. Cells are colored by the expression of Trem2. **(F)** BP enrichment of microglial cells in neonatal SCI mice with down-regulated genes (P<0.05; logFC<0.5), The volcano plot shows the significance of each gene set from the selected library versus its odds ratio. **(G)** 17 hub genes expression of reactive microglia in in neonatal SCI mcie.

Taken together, our findings suggest that adult SCI is characterized by a decrease in innate microglia with reduced cell migration. In contrast, neonatal mice with SCI undergo scar-free healing through the enhancement of innate microglial cell proportion and migration, Notably, TGFβ signal pathway is a potential therapeutic option for promoting SCI recovery, through the activation of microglial cell migration.

### Reactive microglia contribute to the neuroinflammatory response and positive regulate the neural cell death in SCI

We employed gene signatures from reactive microglial cells to conduct a biological process enrichment analysis. Our results indicate that the “Neuroinflammatory Response” is the most significant biological process, and the “Positive Regulation of Neuronal Cell Death” is also a noteworthy biological process associated with reactive microglial cells (Fig. 3A). Furthermore, 17 genes (Trem2, Grn, C1qa, Ccl3, Tyrobp, Apoe, Cst7, C1qb, C1qc, Ccl4, Ch25h, Ctsb, Ctsd, Ctsl, Ctsz, Cxcl13, Egr1) rank in the top ten for biological processes, with Term2 being the foremost among them (Fig. 3B). The analysis of these 17 genes was extended through the utilization of the STRING database to construct a PPI network, demonstrating their close interconnectedness (Fig. 3C). Furthermore, we applied the GSVA method to compute the BP scores for ‘Neuroinflammatory Response’ and ‘Positive Regulation of Neuron Death,’ along with assessing the gene expression levels of Trem2 and neuronal markers (Map2 and Snap25) in ST data. Our analysis revealed a significant enhancement in the ‘Neuroinflammatory Response,’ ‘Positive Regulation of Neuron Death,’ and Trem2 expression within the SCI lesion area. In contrast, The expression of neuronal markers is diminished in the lesion area of SCI, potentially as a result of ‘Positive Regulation of Neuron Death’ by reactive microglia (Fig. 3D). Concurrently, a notable reduction in Trem2 expression was observed over the timeline in neonatal SCI mice exhibiting scar-free healing (Fig. 3E). In addition, the inflammatory response exhibited diminishing trends in neonatal SCI mice (Fig. 3F). Furthermore, the hub genes associated with reactive microglial cells displayed a gradual decrease over the course of the timeline (Fig. 3G). Hence, our belief is that reactive microglia play a pivotal role in driving the neuroinflammatory response in SCI. Consequently, targeting the neuroinflammatory response and Trem2 inhibition could offer a highly promising approach for treating SCI.

### Trem2 inhibition down-regulates neuroinflammatory response and enhances TGFβsignal pathway

Since Trem2 is the key genes of reactive microglia, we performed a comparative analysis of the influence of Trem2, both in the presence and absence of Trem2, on single-cell transcriptional profiles of microglial cells during the pathogenesis of SCI. We utilized publicly available data from the GEO dataset GSE198852(Gao et al., 2023). After completing quality control and filtering, UMAP-based cell clustering effectively distinguished innate and reactive microglial cell clusters based on the naïve and SCI conditions (Fig. 4A). The scatter plot revealed that Tgfbr1 expression was predominantly concentrated within the innate microglial cell cluster, while Trem2 expression was primarily observed in the reactive microglia cluster (Fig. 4B). Next, we isolated the SCI populations and segregated them into two groups based on whether Trem2 was knocked out or not (Fig. 4C, 4D). Interestingly, the expression of reactive microglial gene signatures was notably reduced with Trem2 knockout, particularly among the 17 BP hub genes of reactive microglial cells (Supplementary Fig. S2A, left; Fig. 4E). We further conducted differential gene expression analysis (DEGs) between the groups with and without Trem2 knockout, revealing a total of 264 DEGs (|log2FC | > 0.25; P < 0.05), with 119 of them showing down-regulation in the Trem2 knockout group (Fig. 4F). Subsequently, we performed functional annotation analysis of the down-regulated DEGs, which indicated a reduction in the neuroinflammatory response (Fig. 4G). Additionally, in the WikiPathway signaling pathway database, we observed a decrease in the spinal cord injury and an enhancement in the TGFβ signaling pathway (Fig. 4H, 4I). In light of this, we identified the innate microglia gene signature, along with the expression of 17 hub BP genes. Interestingly, our findings revealed that all these genes were upregulated in the Trem2 knockout model, suggesting that inhibiting Trem2 can effectively transition reactive microglia back to their innate state (Supplementary Fig. S2A, right; Fig, 4J). This is primarily attributed to the robust interaction between TREM2 (PDB: 5UD7) and TGFBR1 (PDB: 5E8S), characterized by their competitive intermolecular interactions involving hydrogen bond propensities and van der Waals forces, as demonstrated in the Chimera 3D Model (Fig. 4K, left). In the molecular surface view depicted in space-filled docking models (Fig. 4K, right), the strength of interaction between these two proteins is prominently evident. Thus, we conclude that targeting Trem2 can effectively down-regulate the neoroinflammatory response and enhance the TGFβ signal pathway by modulating the gene signature of reactive microglia.

**Fig. 4.**
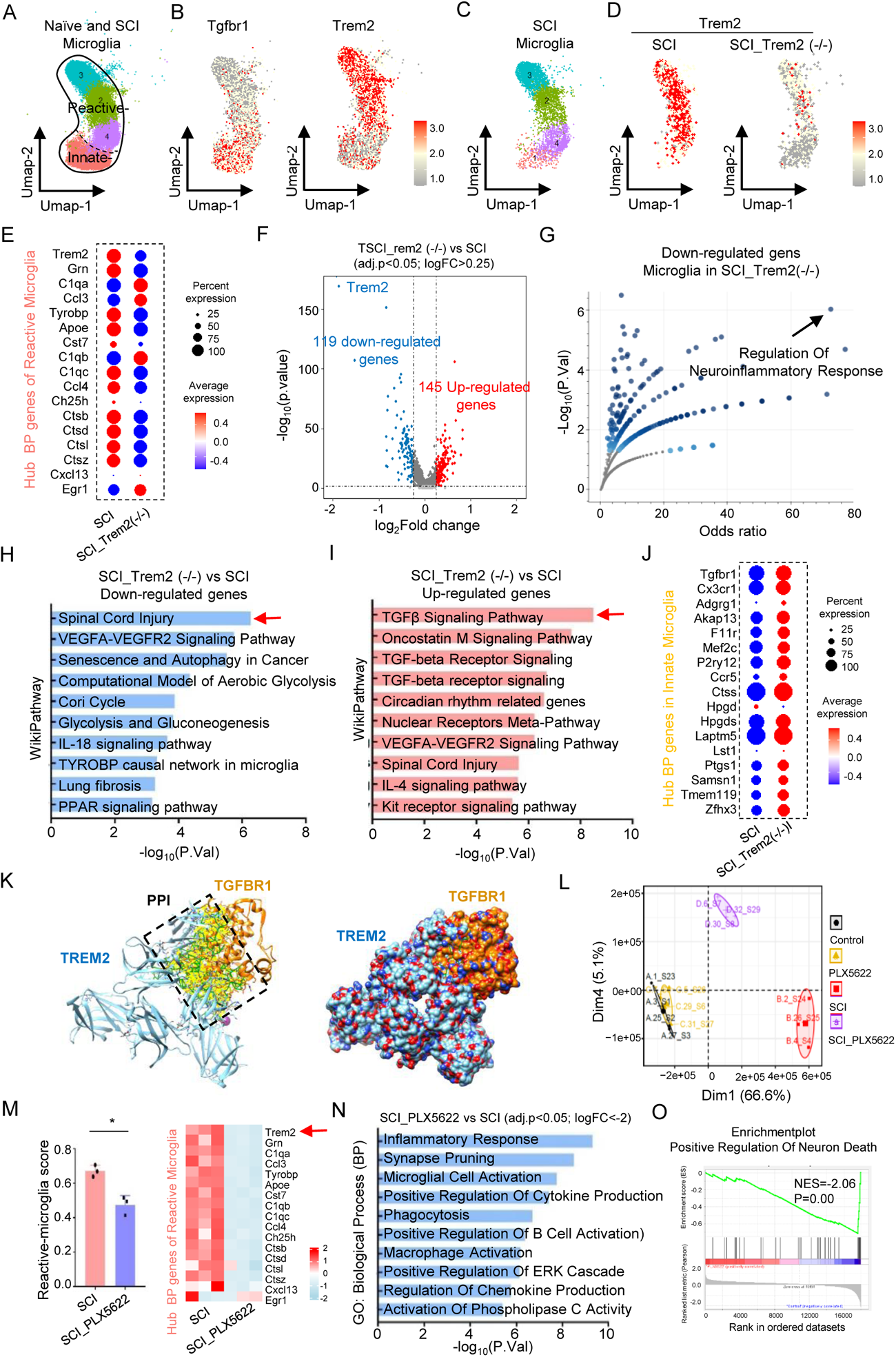
Trem2 inhibition down-regulates neuroinflammatory response and enhances TGFβ signal pathway. **(A)** Single-cell RNA sequencing of microglial cells in spinal cords under naïve and SCI conditions. We analyzed the resulting clusters with the UMAP method and differentiated cell populations by color (GSE198852). **(B)** Scatter plot show the expression of Tgfbr1 and Trem2. (**C**) Single-cell RNA sequencing of microglial cells in spinal cords from SCI and SCI_Trem2-/-mice. **(D)** Trem2 expression in SCI and SCI_Trem2-/-mice. **(E)** Hub genes of Biological Process (BP) in reactive microglia between SCI and SCI_Trem2-/-mice. **(F)** Volcano plot show the different genes between SCI and SCI_Trem2-/-mice (adj.p<0.05; logFC>0.25). **(G)** Microglial cell enrichment in T SCI_Trem2-/-mice, with down-regulated genes highlighted (P<0.05; logFC>0.5), visualized via Appyter. The volcano plot showcases gene set significance against its odds ratio. **(H)** Enriched signal pathway in microglial cells from Trem2(-/-) SCI, highlighting down-regulated genes, **(I)** and highlighting upregulated genes. **(J)** Hub BP genes of innate microglia between SCI and SCI_Trem2-/-mice. **(K)** The Chimera 3D Model illustrates the interaction between TREM2 (PDB: 5UD7) and TGFBR1 (PDB: 5E8S), highlighting their competitive intermolecular interactions, including hydrogen bond propensities (green lines) and van der Waals forces (yellow lines) on the left. On the right, the molecular surface view presents TREM2 and TGFBR1 as space-filled models, distinguished by heteroatom coloring. **(L)** Principal component analysis (PCA) biplot of transcriptional profiles with 95% confidence ellipsoids for Control (uninjured), SCI, PLX5622, and SCI_PLX5622 (GSE196928) panels. **(M)** Gene Set Variation Analysis (GSVA) by using reactive microglia gene signatures between PLX5622+SCI and SCI (left). Hub BP genes of reactive microglia in PLX5622+SCI vs. SCI (right). **(N)** Enrichment of BP using down-regulated genes between PLX5622+SCI and SCI (adj.p<0.05; logFC<-2). **(O)** The Gene Set Enrichment Analysis (GSEA) enrichment plot of Positive Regulation Of Neuron Death in PLX5622+SCI vs. SCI.

While specific inhibitors for Trem2 are currently lacking, some indirect agents, such as PLX5622 (Yun Li et al., 2020), have already demonstrated promising effects. Therefore, we obtained data on the use of pharmacologically depleted microglia (PLX5622) and used bulk RNA sequencing to reveal the cellular and molecular responses to SCI controlled by microglia. Principal Component Analysis (PCA) initially unveiled correlations among panels of control (uninjured), SCI, PLX5622, and SCI_PLX5622. This analysis revealed that the majority of control and PLX5622 cells clustered together in the same principal component, while SCI and SCI_PLX5622 exhibited a distinct component. This observation illustrates that PLX5622 has little impact on the normal spinal cord but significantly influences the SCI component (Fig. 4L). A comparison of transcriptional profiles between SCI_PLX5622 and SCI revealed a significant decrease in microglia gene signatures with PLX5622 treatment (Supplementary Fig. S2B). Furthermore, the final assessment of the reactive microglia gene signature GSVA score, particularly the 17 hub BP genes, with a notable emphasis on Trem2, also revealed a significant decrease following PLX5622 treatment (Fig. 4M). The most noteworthy biological process that exhibited a decrease was the inflammatory response, coupled with the inhibition of neuron death, as revealed through GSEA analysis (Fig. 4N, 4O).

### iPSC-derived microglia have the potential to replace the functions of naive microglial cells to combat SCI

As mentioned earlier, the replacement of innate microglial cells with reactive microglia cells positive regulate the neuronal cell death in SCI. Therefore, inhibiting the reactive microglia and subsequently transplanting innate microglial cells could represent a promising strategy for combating SCI. To test this hypothesis, we performed a comprehensive data reanalysis using three efficient microglia replacement strategies: bone marrow transplantation (mrBMT), peripheral blood (mrPB), and microglia transplantation (mrMT) (GSE155499) (Xu et al., 2020). UMAP analysis unveils that the bulk of mrMT cells grouped with naive microglia cells, whereas, in contrast, the majority of mrBMT and mrPB cells resided in distinct clusters (Fig. 5A). Innate microglia gene signatures, particularly the 17 hub gens exhibit similarities between mrMT and naive microglia cells but display lower expression levels in mrBMT and mrPB cells (Supplementary Fig. S3; Fig. 5B). The mrMT and microglia cell cluster demonstrated a higher capacity for cell migration compared to other groups (Fig. 5C). The results demonstrate that microglia transplantation effectively fulfills the role of innate microglia, potentially enhancing SCI recovery. Nonetheless, the restricted availability of donor cells for microglia transplantation could potentially hinder its widespread adoption in future SCI treatments. To overcome this limitation, induced pluripotent stem cells (iPSC) dedrived microglia can be generated from somatic cells, and their innovative development has led to rapid progress in the field of SCI regeneration (Nagoshi & Okano, 2018). We will explore the data from GSE89189 to compare the transcriptome profiling of iPSC derived microglia with that of human primary fetal microglia and adult microglia (Abud et al., 2017).

**Fig. 5.**
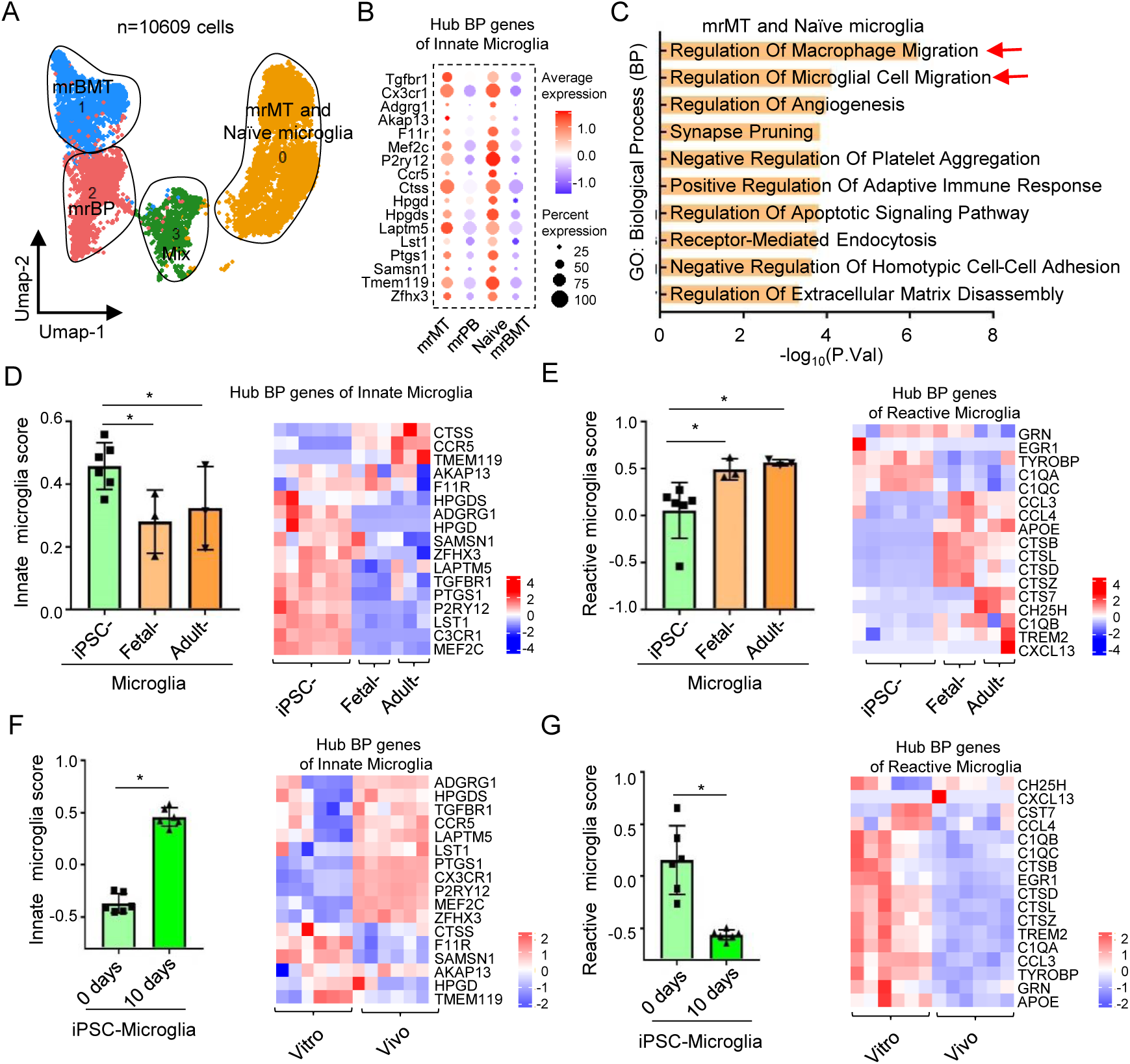
iPSC derived microglial cells transplantation is a promising approach to combat SCI. **(A)** Single-cell RNA-seq for naive microglia and bone marrow transplantation (mrBMT), peripheral blood (mrPB), and microglia transplantation (mrMT) (GSE155499). **(B)** Hub BP genes of innate microglia among naive microglia, mrBMT, mrPB and mrMT. **(C)** Biological Process (BP) enrichment of mrMT and naïve microglia (adj.p<0.01; logFC>0.5). **(D)** Gene Set Variation Analysis (GSVA) by using innate microglia gene signatures (left), Hub BP genes of innate microglia among naive microglia, mrBMT, mrPB and mrMT (right). **(E)** GSVA by using reactive microglia gene signatures (left), Hub BP genes of reactive microglia among naive microglia, mrBMT, mrPB and mrMT (right). **(F)** Gene Set Variation Analysis (GSVA) by using innate microglia gene signatures (left), Hub BP genes of innate microglia between 0 and 10 days after transpnaltation (right). **(G)** Gene Set Variation Analysis (GSVA) by using reactive microglia gene signatures (left), Hub BP genes of reactive microglia between 0 and 10 days after transpnaltation(right).

Next, we apply GSVA analysis to calculate the innate- and reactive-microglia score by using their own gene signatures. Our research has revealed that iPSC-derived microglia exhibit significantly elevated innate microglia scores. Additionally, the 17 BP hub genes exhibit a consistent trend in line with their GSVA score (Fig. 5D). However, when it comes to reactive microglia scores and their corresponding 17 BP hub genes, they appear to be lower in comparison to fetal and adult microglia (Fig. 5E). This show that iPSC-derived microglia outperform the naïve microglia. To assess the long-term persistence of iPSC-derived microglia in vivo, we sought additional data from the GEO datasets (GSE139192) (Svoboda et al., 2019). Remarkably, iPSC-derived microglia in the CNS after 10 days exhibit significantly higher expression of the innate microglia gene signature, with a noticeable trend in majority of the 17 hub genes (Fig. 5F). Conversely, they demonstrate a decreased expression of the reactive microglia gene signature, also displaying a discernible trend in almost the 17 hub genes (Fig. 5G). This show that iPSC-derived microglia performed better in vivo. Taken together, iPSC-derived microglia have the potential to replace the functions of naive microglial cells, and they perform even more effectively in the in vivo CNS. This strongly reinforces our belief that this approach holds great promise in combating SCI.

## Disscussion

Microglia cells display diverse multi-cellular signal responses and biological processes that can either contribute to secondary injury or facilitate neural repair following SCI, in response to various internal and external factors (Wang et al., 2022). The present research demonstrates a strong correlation between microglial cells and SCI recovery, where a decrease in innate microglia, accompanied by an increase in reactive microglia, are observed. Furthermore, TGFβ signaling pathway regulate innate microglial cell migration to improve SCI recovery however, reactive microglial cell contribute to the neuroinflammatory response can positive regulate the neural cell death in SCI. Specifically, inhibiting Trem2 in reactive microglial cells not only reduces the inflammatory response but also mitigates spinal cord injury, while enhancing the TGFβ signaling pathway shows promise. Another promising avenue involves the utilization of iPSC-derived microglial cells, which have demonstrated their ability to enhance the potential for replacing the functions of naive microglial cells, performing even more effectively in the in vivo CNS. This strongly reinforces our belief that inhibiting reactive microglial cells through Trem2 and subsequently transplanting iPSC-derived microglial cells with TGFβ modulation could prove to be a promising approach in treating SCI.

Over the past few decades, microglial biological process has received relatively more attention. While there is no clear consensus on whether it is beneficial or detrimental in spinal cord repair. Some researchers support the beneficial functions of microglia, because efficient clearance of tissue debris is critical in reconstructing and reorganizing neuronal networking following neural system injuries (Fu et al., 2014; Sierra et al., 2010; Weldon et al., 1998). However, excessive reactive microglial phagocytosis can have detrimental effects on spinal cord repair as it can lead to the removal of injured neurons that still have some function, while also increasing the inflammatory response and disrupting homeostasis of neuron survival. In the uninjured central nervous system (CNS), microglia exhibit high migratory activity during the perinatal period of development due to their innate nature (Dibaj et al., 2010; Kettenmann et al., 2011). However, after an adult sustains a SCI, glial scarring suppresses the migration of microglia over long distances, inhibiting their accumulation at the site of damage (Yi Li et al., 2020). Even more concerning, the activation of TREM2 in neurotraumatic injuries, such as traumatic brain injury and peripheral nerve injury, can lead to adverse effects, including exacerbated neuronal loss (Kobayashi et al., 2016; Saber et al., 2017). Our findings also corroborate this observation in the context of SCI. Therefore, the development of a novel approach to modulate the intrinsic properties and reactive functions of microglial cells holds immense promise. For instance, TGFβ has demonstrated the potential to enhance microglial cell migration, while TREM2 inhibition or PLX5622 can effectively suppress the inflammatory response in reactive microglial cells in our study. Previous studies have reported that microglia depend on TGF-β signaling, which sheds light on microglial biology and suggests the potential for targeting microglia as a treatment for SCI (Butovsky et al., 2014; Lodge & Sriram, 1996). Notably, the most significant improvements in locomotor function in mice were observed when PLX5622 chow was administered two to three weeks prior to SCI and continuously thereafter (Bellver-Landete et al., 2019; Brennan et al., 2018). TREM2-ablated microglia displayed a substantial decrease in lesion size and a noteworthy enhancement in locomotor function after SCI, underscoring the crucial implications for the development of TREM2-targeting therapeutics (Gao et al., 2023).

Recent advances in the generation of microglia from iPSC have provided exciting new approaches to examine and decipher the biology of microglia. By promoting the survival and regeneration of neurons and regulating neurotransmission, these cells have the potential to improve motor function and sensory outcomes in individuals with neurological diseases (Kang & Guo, 2022; Khazaei et al., 2014; Li et al., 2021; Nishida et al., 2020). For instance, when hiPSC-derived microglia are transplanted into the neonatal mouse brain, they adopt a phenotype and gene expression signature reminiscent of resting microglia found in the human brain (Svoboda et al., 2019). Another study demonstrated that human iPSC-derived microglia cells integrated into the mouse retina and faithfully replicated the characteristics of native microglia cells (Ma et al., 2023). While there is no direct evidence reporting the benefits of iPSC-derived microglia transplantation for motor and sensory recovery in SCI, there is a research report indicating that the transplantation of microglia with anti-inflammatory effects promotes the recovery of motor function after SCI in mice (Kobashi et al., 2020). Drawing from our findings, iPSC-derived microglia have demonstrated the potential to augment the replacement of native microglial cell functions. This suggests the promising prospect of differentiating iPSCs into microglial cells in the near future as a viable strategy for treating SCI.

Nonetheless, it is crucial to acknowledge that the study we have reviewed is confined to bioinformatics analysis, and there are several substantial obstacles that must be surmounted to conduct a successful in vitro and in vivo study. In this particular study, the most promising aspect lies in the efficient clearance of microglia in the CNS using drug-specific inhibition of Trem2, which avoids significant side effects. Subsequently, iPSC-derived microglial cell transplantation was performed in the injured mouse spinal cord. Following this, transplanted cells were selectively stimulated daily through TGFβ injections, resulting in promising improvements in motor functional recovery. This represents an area of future work that truly warrants our dedicated efforts.

## Conclusions

In summary, as depicted in Fig. 6, in the adult SCI model, innate microglial cells are replaced by reactive microglia. Alongside these changes come alterations in their associated biological processes and signal pathways, such as decreased TGF-regulated cell migration and heightened Trem2-mediated neuroinflammatory response, which can impede SCI recovery. However, the scenario is different in neonatal mice with SCI, where a scar-free healing of the injured spinal cord is observed. Therefore, inhibiting Trem2-mediated neuroinflammatory responses and transplanting iPSC-derived microglia with long-term TGFβ stimulation may offer potential improvements in SCI recovery.

**Fig. 6.**
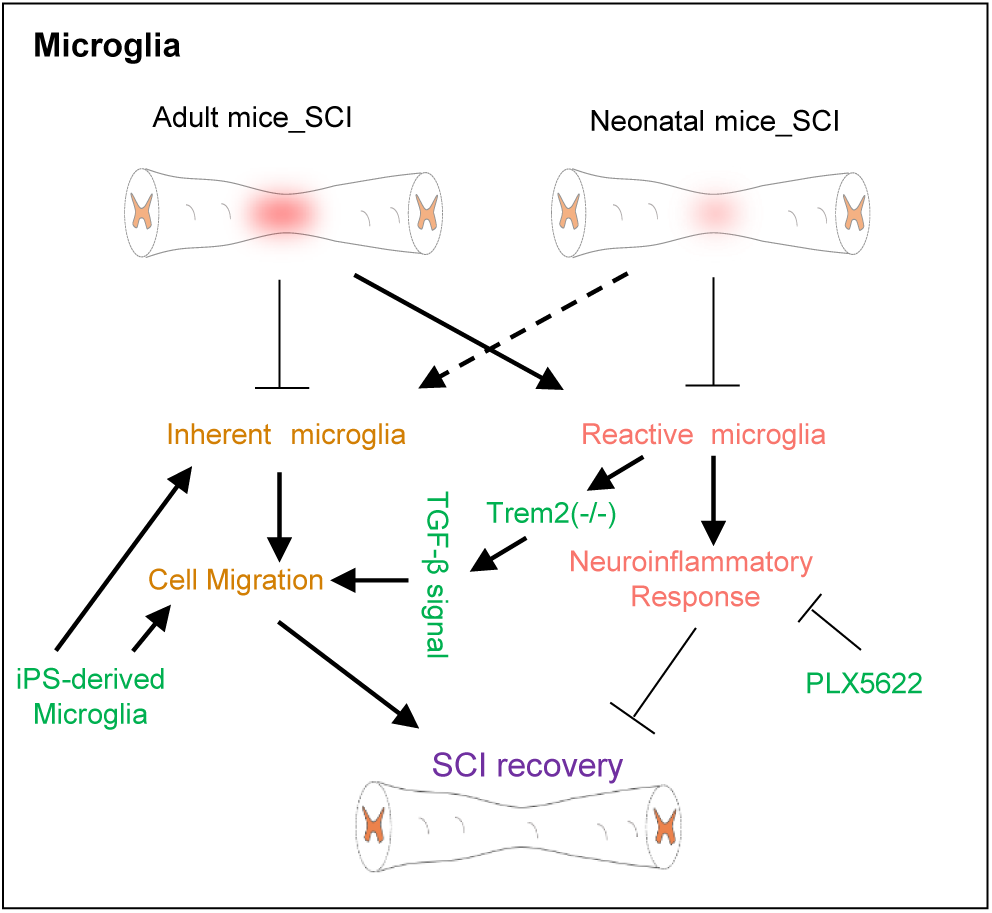
Overview of the signal pathways and biological functions in chondrosarcoma. In adult spinal cord injury (SCI) models, innate microglial cells transform into reactive microglia. This shift leads to changes in biological processes and signaling pathways, including reduced TGF-regulated cell migration and increased Trem2-mediated neuroinflammatory responses, which can hinder SCI recovery. Conversely, neonatal mice with SCI show scar-free spinal cord healing. Consequently, strategies like inhibiting Trem2-mediated neuroinflammatory responses and transplanting iPSC-derived microglia with prolonged TGFβ stimulation may hold promise for enhancing SCI recovery.

## Abbreviations

Adgrg1: Adhesion G Protein-Coupled Receptor G1
Akap13: A-Kinase Anchoring Protein 13
Apoe: Apolipoprotein E
C1qa: Complement C1q A Chain
C1qb: Complement C1q B Chain
C1qc: Complement C1q C Chain
Ccl3: C-C Motif Chemokine Ligand 3
Ccl4: C-C Motif Chemokine Ligand 4
Ccr5: C-C Chemokine Receptor Type 5
Ch25h: Cholesterol 25-Hydroxylase
Cst7: Cystatin F
Ctsb: Cathepsin B
Ctsd: Cathepsin D
Ctsl: Cathepsin L
Ctss: Cathepsin S
Ctsz: Cathepsin Z
Cx3cr1: C-X3-C Motif Chemokine Receptor 1
Cxcl13: C-X-C Motif Chemokine Ligand 13
Egr1: Early Growth Response 1
F11r: F11 Receptor
Grn: Granulin
Hpgd: Hydroxyprostaglandin Dehydrogenase 15-(NAD)
Hpgds: Hematopoietic Prostaglandin D Synthase
iPSCs: induced pluripotent stem cells
Laptm5: Lysosomal Protein Transmembrane 5
Lst1: Leukocyte Specific Transcript 1
Map2: microtubule-associated protein 2
Mef2c: Myocyte Enhancer Factor 2C
P2ry12: Purinergic Receptor P2Y12
Ptgs1: Prostaglandin-Endoperoxide Synthase 1
Samsn1: SAM Domain-Containing Protein Samsn1
SCI: Spinal cord injury
Snap25: Synaptosome associated protein 25
Tgfbr1: Transforming growth factor beta receptor 1
TGFβ: Transforming growth factor beta
Tmem119: Transmembrane Protein 119
Trem2: Triggering receptor expressed on myeloid cells 2
Tyrobp: TYRO Protein Tyrosine Kinase Binding Protein
UMAP: Uniform Manifold Approximation and Projection
Zfhx3: Zinc Finger Homeobox 3

## Acknowledgments

We are very grateful to the participating universities and China Scholarship Council.

## Author contributions

Y.S. conceived the study and generated Fig.s and wrote the manuscript, Q.Z. collected and analysed the data, Q.F. and J. L. performed a part of data analysis. C. Z. and W. L help to generated Figures. All authors reviewed the manuscript.

## Availability of data and materials

All data generated or analyzed during this study have been incorporated into this published article and its accompanying supplementary information.

## Declarations

### Ethics approval and consent to participate

Not applicable.

### Consent for publication

Not applicable.

---

**Supplementary Fig. S1**. The heatmap illustrates the expression of gene signatures for innate (A) and reactive (B) microglia in neonatal SCI mice.

**Supplementary Fig. S2.** (A) Dot plot demonstrates the expression of reactive and innate microglia gene signatures between SCI and SCI_Trem2(-/-) SCI mice. (B) Heatmap depicts the expression of reactive microglia gene signatures between SCI_PLX5622 and SCI mice.

**Supplementary Fig. S3.** Expression of innate microglia gene signatures across naive microglia, mrBMT, mrPB and mrMT

**Supplementary Table S1.** Distinct sets of genes of each cluster in single cell data.

